# A computational model of MGUS progression to Multiple Myeloma identifies optimum screening strategies and their effects on mortality

**DOI:** 10.1101/208645

**Authors:** Philipp M. Altrock, Jeremy Ferlic, Tobias Galla, Michael H. Tomasson, Franziska Michor

## Abstract

Recent advances uncovered therapeutic interventions that might reduce the risk of progression of premalignant diagnoses, such as from Monoclonal Gammopathy of Undetermined Significance (MGUS) to multiple myeloma (MM). It remains unclear how to best screen populations at risk and how to evaluate the ability of these interventions to reduce disease prevalence and mortality at the population level. To address these questions, we developed a computational modeling framework. We used individual-based computational modeling of MGUS incidence and progression across a population of diverse individuals, to determine best screening strategies in terms of screening start, intervals, and risk-group specificity. Inputs were life tables, MGUS incidence and baseline MM survival. We measured MM-specific mortality and MM prevalence following MGUS detection from simulations and mathematical precition modeling. We showed that our framework is applicable to a wide spectrum of screening and intervention scenarios, including variation of the baseline MGUS to MM progression rate and evolving MGUS, in which progression increases over time. Given the currently available progression risk-point estimate of 61% risk, starting screening at age 55 and follow-up screening every 6yrs reduced total MM prevalence by 19%. The same reduction could be achieved with starting age 65 and follow-up every 2yrs. A 40% progression risk reduction per MGUS patient per year would reduce MM-specific mortality by 40%. Generally, age of screening onset and frequency impact disease prevalence, progression risk reduction impacts both prevalence and disease-specific mortality, and screeenign would generally be favorable in high-risk individuals. Screening efforts should focus on specifically identified groups of high lifetime risk of MGUS, for which screening benefits can be significant. Screening low-risk MGUS individuals would require improved preventions.

## INTRODUCTION

Multiple myeloma (MM) is the second most common hematologic malignancy in the US, representing 1. 8% of new cancer cases and 2.1% of cancer deaths annually (1). MM is an incurable plasma-cell malignancy (2). Patients show abnormal levels of the paraprotein M-protein (3), indicating a monoclonal cell population and end organ damage such as lytic bone lesions (4). Almost all MM patients progress from a precursor condition called Monoclonal Gammopathy of Undetermined Significance (MGUS), displaying only M-protein spikes (4). The MGUS condition exists in approximately 3% of the population of age 50 or higher (5), and men show higher age-adjusted incidence rates than women (6).

Recent advances suggest that the rate of progression to MM can be altered by therapeutic interventions (7, 8). For example, obesity—a modifiable risk factor for MM—is associated with increased risk (9–11). Furthermore, metformin is associated with a reduced progression of MGUS to MM, potentially delaying MM by four years in type-2 diabetics with MGUS (8). Reduced risk is also associated with regular use of aspirin (7). Although causal relationships of these associations, as well as molecular mechanisms, are uncertain, these findings suggest that pharmacological and other interventions have the potential to reduce the risk of progression from MGUS to MM. It is therefore of particular interest to investigate the effects of screenings of the general population, or of specific subpopulations and their distribution across risk groups, with the goal of early MGUS detection and maximal potential reduction of progression risk to MM.

Here we designed a computational model that describes incidence of MGUS and progression to MM, specific MGUS screening scenarios, and potential epidemiological changes, implemented after detection. Our model is based on life tables and epidemiological data of MGUS and MM, which depend on genetic background, gender and age (12, 13) and correlate with ethnicity (14). Using simulations and analytical results, we assessed whether a given reduction in the progression risk after a positive MGUS screen can reduce MM prevalence and lead to changes in MM-specific mortality (or survival). Our work can be used to identify optimal screening strategies and can assess the utility of interventions targeting MM precursor states.

**Table 1:** Important parameters used for computational and mathematical modeling.

| Parameter | Description | Range or Value | References |
| --- | --- | --- | --- |
| $a$ | Age | 0-100 years | (17) |
| $d(a)$ | Probability to die of any cause at age $a$ | 0-1. Age dependent | (1, 17) |
| $d_{MM}$ | Probability to die of MM, (see Supplement) | 0.1295 per MM patient per year | (36, 37) |
| $m(a)$ | Incidence rate of MGUS | 0-1 per person per year, age dependent, risk-group dependent | (12), this work |
| $p$ | Probability of progression from MGUS to MM | 0-0.15 per person per year, depending on progression model, disease evolution | (22, 23, 28), this work |
| $a_0$ | Age of first MGUS screen | 20-50 years | This work |
| $\Delta a$ | Interval between screens | 1-15 years | This work |
| $r$ | Reduction in progression, conditional on MGUS detection | 0-1. For example, if $r=0.5$, then $p=0.5 \times 0.01=0.005$ per MGUS patient per year | (7, 8) |

## MATERIALS AND METHODS

We developed a Markov chain model (**Fig. 1A**) in which healthy individuals transition to an undetected MGUS stage, from which they can transition to detected MGUS if screened. An individual with MGUS progresses to overt MM at a certain rate per year; however, a positive MGUS screening result reduces the rate of progression to MM (**Fig. 1 B, C**). Individuals may die at any point but mortality is larger for those with MM than others. We performed stochastic simulations and derived an analytical framework to assess MM mortality and prevalence reduction after screening (Supplement).

**Figure 1:**
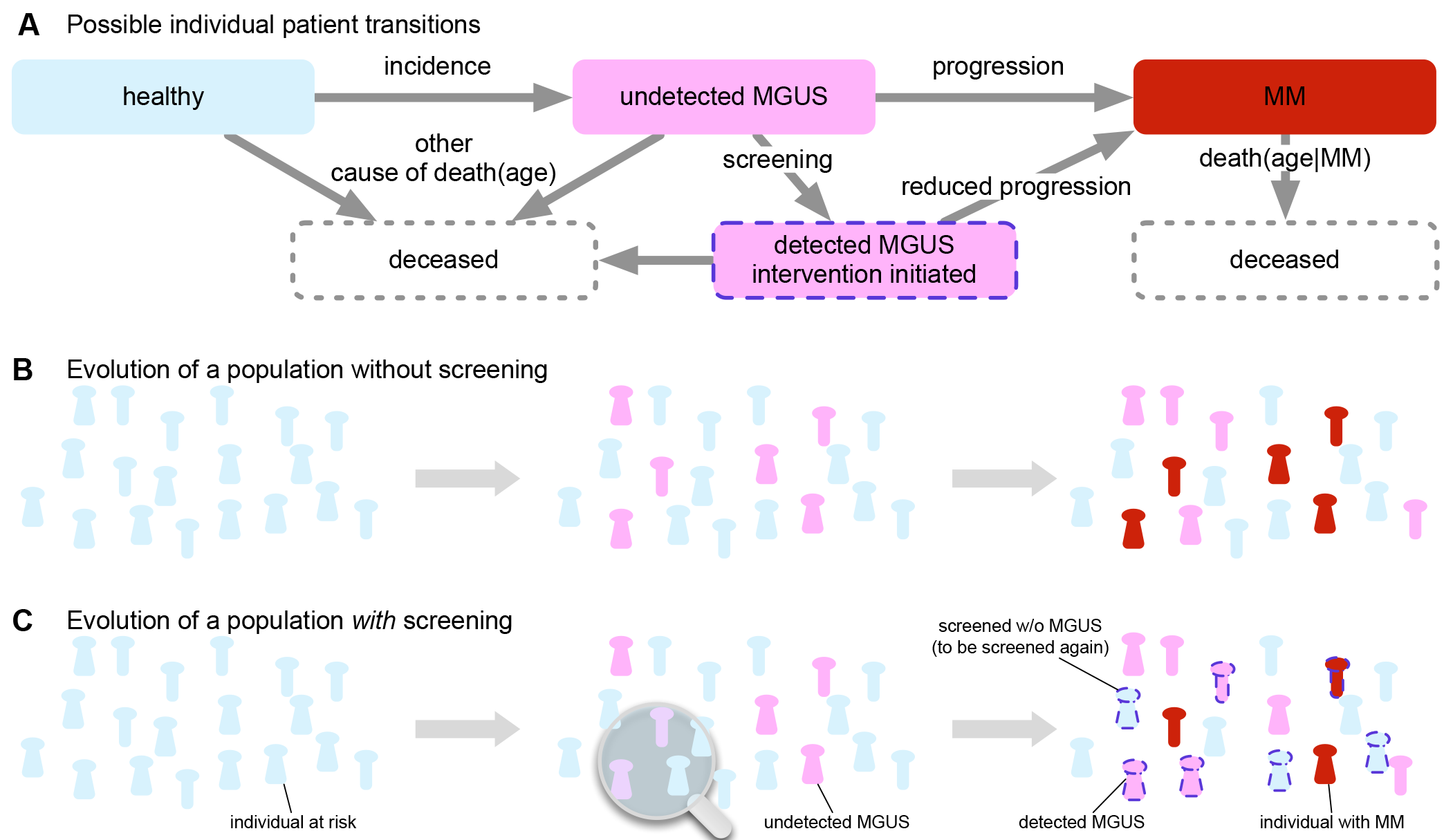
Population dynamics of unscreened and screened MGUS as well as MM individuals. **A**: The individual transitions from healthy to MGUS to MM can be modeled as Markov chain. The transitions describe incidence and screening of Monoclonal Gammopathy of Undetermined Significance (MGUS) and progression to Multiple Myeloma. The four possible states are healthy (light blue), undetected MGUS (pink), detected MGUS (pink with dashed outline), and MM (red). **B**: Example time evolution of a cohort at risk of MGUS and subsequent MM, without screening. Undetected MGUS cases accumulate and can lead to a baseline number of MM cases. **C**: Time evolution of a cohort with screening, and intervention that reduces MGUS to MM progression. MGUS cases accumulate; individuals are screened and receive preventive treatment if positive for MGUS, leading to a lower number of MM cases (red, few screened individuals may develop MM nonetheless).

### Model inputs and outputs

We were interested in screening outcomes in mixture-populations composed of individuals with different MGUS lifetime risks. We distinguished non-African American and African Americans as low-risk (baseline) and high-risk individuals, respectively. From baseline, high-risk individuals carry an average two-fold increase in lifetime risk of MGUS (13, 15). Calculations of the respective MGUS incidence rates are displayed in the Supplement. Further, we used a crude birth rate for the total population and life tables to calculate death events of healthy and MGUS individuals (high-and low-risk males and females), MM-specific death rates, and a fixed MGUS to MM progression rate for unscreened individuals. A screening scenario was specified by three parameters: age of the individual when receiving the first screen, *a*_0_, spacing between follow-up screens, Δ*a*, and risk reduction *r*, after a positive screen (**Table 1**). As model outputs, we were interested in the effects of varying screening scenarios on the MM-specific mortality after MGUS detection and on the fraction of individuals with MM of all ages. We initiated all simulated populations according to the age distribution of the population in the US according to the 2013 census (16, 17), with a fixed fraction of healthy high-risk individuals of 20%. Although the fraction of African-Americans in the US is about 13% (17), we estimated that the genetic diversity in the US would further contribute to high-risk.

### Stochastic model

We simulated the Markov chain model (**Fig. 1A**, Supplement) by using a fixed crude birth rate (18), age-dependent death rates for healthy and MGUS individuals (17), and a fixed death rate for MM patients (19). Eventually, the simulated populations assumed a stationary state independent of the initial age distribution (20) (Fig. S1). From the baseline low-risk MGUS incidence, adapted from (12), we calculated elevated incidence rates per life year for specific risk groups. In our simulations, high-risk African-Americans experience MGUS incidence that exponentially increase over age such that lifetime risk is about 2-fold higher than baseline (low-risk) (13, 21). Progression to MM was mostly constant across risk groups (22) and occurred at a rate of *p*=0.01/year in MGUS-positive but unscreened individuals (23). Screening meant that starting at age *a*_0_, individuals were screened each year with prob. 1/Δ*a*, such that their average time between screens was Δ*a*. Positively screened individuals were assumed to progress at a reduced rate of *r*\**p*. Recent studies have estimated *r*=0.61 for regular aspirin users (7). From simulations, individual ages, MGUS status, MGUS screening and MM status were recorded (Supplement). This approach allowed us to calculate MGUS and MM prevalence, distribution of MM age at diagnosis, and MM-specific mortality. We also devised a model to calculate MGUS and MM prevalence and mortality analytically (Supplement). Using this framework we calculated the fractions of MGUS individuals, *M*, at a specific age for any risk group, the fraction of MM individuals proportional to *M*, as well as the MM-specific mortality for a given number of years after MGUS detection.

## RESULTS AND DISCUSSION

### Prevalence of MM when screening for MGUS

We performed stochastic simulations of our agent-based model to investigate the effects of different conditions on MGUS and MM prevalence and mortality. As expected, the proportions of individuals with MGUS and MM varied with the fraction of high-risk persons in the population (**Fig. S1**). An increasing risk reduction after a positive MGUS screen drastically diminished the fraction of MM cases while increasing the fraction of MGUS cases (**Fig. 2A**). To validate our results we compared our findings to those of Birmann *et al.* (7), where in a cohort of 163,810 men and women, 82 individuals were associated with the baseline progression risk and 44 were associated with the lowest progression risk measured, with a value of *r*=0.61, in long-term aspirin users (95% CI between 0.41 and 0.95) (7). These authors reported a reduction of 40% in MM cases linked to aspirin use. Based on this study we estimated a reduced risk in progression from MGUS to MM of *r*=0.61 (point estimate). For this value our predictions of about 60% lie in Birmann *et al.*’s confidence interval for *r*.

Changes in onset age of screening, *a*_0_, and spacing, Δ*a*, impacted MM risk reduction similarly (Fig. 2B, **Table S4**). For example, for a fixed *r*=0.61, *a*_0_=45y and Δ*a*=8y reduced MM prevalence to 77.2%, whereas *a*_0_=65 and Δ*a*=8y reduced MM prevalence to 78.6% relative to the *r*=1, respectively (Table S4). Even for nearly complete risk reduction (*r* close to 0) and rare screening (Δ*a*=8y), *a*_0_=45y reduced MM cases by 60% (Fig. S2), and *a*_0_=65y by about 38%. **Figs. 2C,F** shows the impact of Δ*a* and *a*_0_ on the age distribution of MM diagnoses, varying *r*. These plots show normalized violin plots give the probability to find an individual of a specific age with MM in our simulations. The bottleneck near *a*_0_, was clearly more pronounced for lower values of *r*. Hence, both the number of MM cases and MM diagnosis age are very sensitive to changes in the progression risk, screening interval and screening start age.

**Figure 2:**
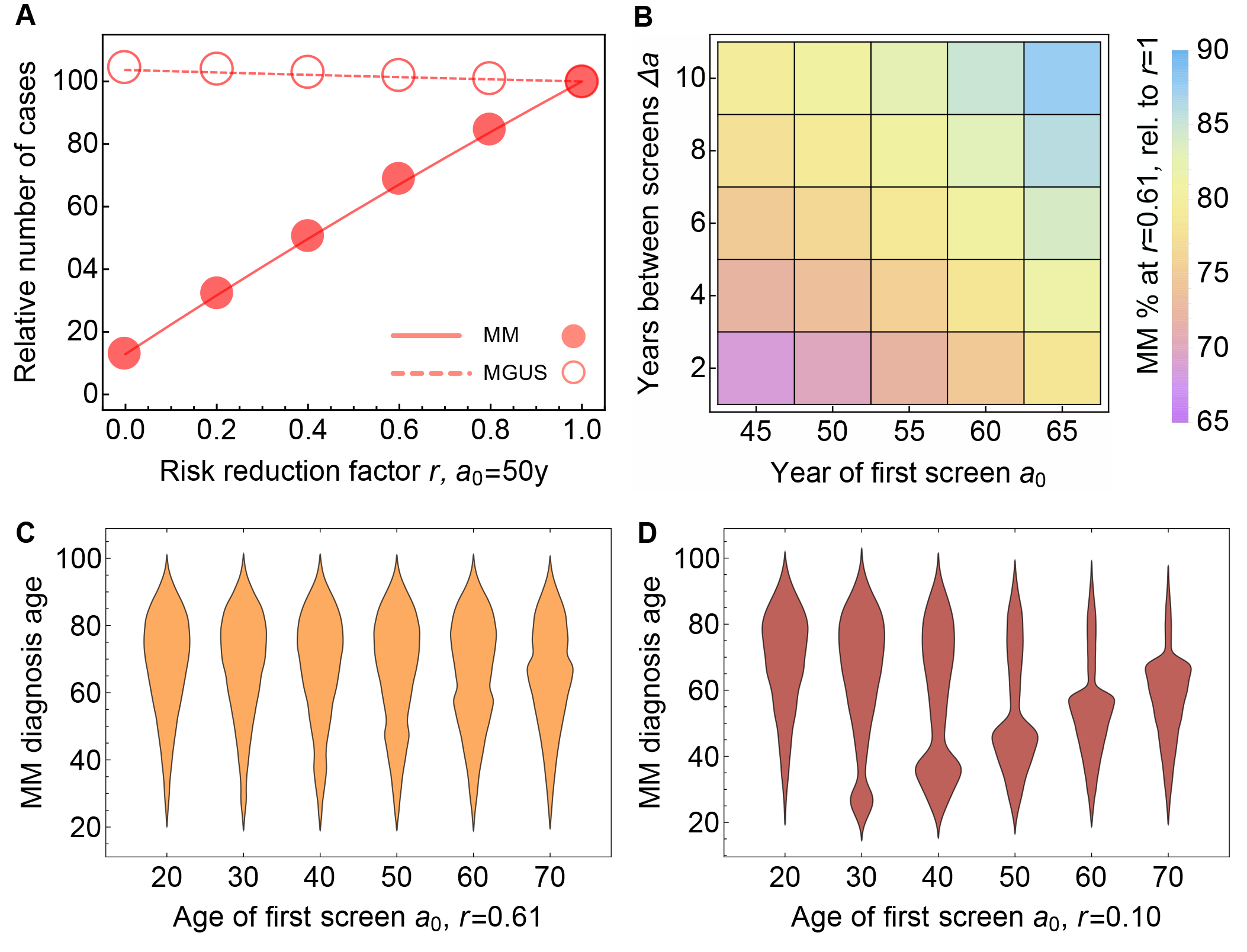
Number of MM-cases, age at MM diagnosis and variability of screening strategy. **A**: When MGUS screening was applied we measured the change in the fractions of MGUS (dashed line, circles) and MM (solid line, disks) with respect to changing the risk reduction factor *r* (symbols: simulations, lines: analytical model, see Supplement and also Figure S2), *a*_0_=50y, Δ*a*=1y, with variable r. With *r*=0.61, the MM fraction dropped below 70% of its value at *r*=1 (where screening had no effect on progression). **B**: Variability in MM fraction at *r*=0.61, with respect to changes in *a*_0_ and *Δa* (analytical approach, point estimates, see Table S4). **C, D**: Distributions of age at MM diagnosis (Δ*a*=1y), varying *a*_0_, fixed *r*=0.61 (**E**), *r*=0.1 (**F**). Width in these violin plots is equal to probability of MM diagnosis at that age. All point estimates were calculated from a simulation of about 10^8^ individuals.

### Lead-time bias and cumulative MM-specific mortality

Screening can cause a lead-time bias in that the survival time after a positive MGUS screening outcome is typically longer than the survival time after direct clinical presentation of MM, with or without screening; the difference between these two times is the lead-time bias (24, 25). Because lead-time bias overshadows actual survival benefits of screening in clinical settings where this time difference may not be directly observed, disease-specific mortality is more appropriate (26). We determined the expected lead-time bias by a comparison of survival in unscreened (control) and screened population simulations (**Fig. 3A**). Median survival post MM diagnosis in the control group was 4-5 years. Median survival post MGUS detection (*a*_0_=50y, *Δa*=1) was 15 years for *r*=1.0 (and similar for *r*=0.61) and 17 years for *r*=0.1. Thus the lead-time bias here would be 10 years.

**Figure 3:**
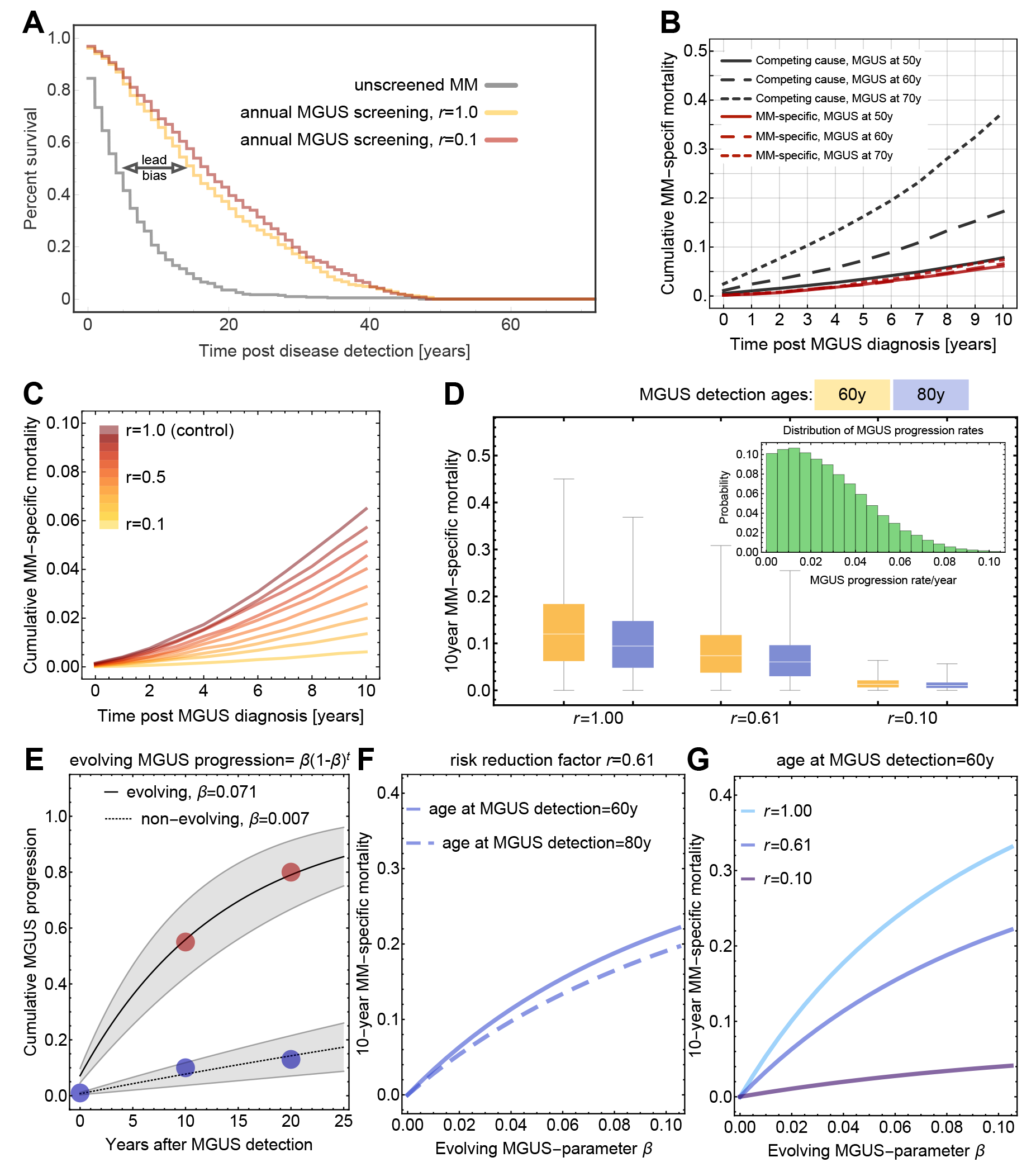
Lead-time bias, cumulative MM-specific mortality, and MGUS to MM progression variability. All simulations were performed with populations of 10^8^ healthy individuals (20% high-risk). **A**: Potential lead-time bias, comparing (i) median survival after MM diagnosis w/o screening (gray: median survival 4 years) and (ii) w/screening (yellow: median survival 15 years, red: median survival 17 years after MGUS screen, respectively). Without screening, disease detection was the event of MM diagnosis. With screening, disease detection was diagnosis of asymptomatic MGUS. **B**: Cumulative MM-specific mortality in years following MGUS detection were measured for the groups of 50, 60, or 70 years of age at MGUS detection, *a*_0_=50, *Δa*=1, and *r*=1. In older patients, death of other cause becomes more dominant. **C**: MM-specific mortality changed dramatically with *r* (*a*_0_=50, *Δa*=1), here shown for individuals diagnosed with MGUS at age 60y, sampled from simulations. **D**: MM-specific mortality is influenced by variability in MGUS to MM progression rate (22) (inset, truncated Normal distr.; mean 0.01, standard dev. 0.03) on, for different *r*, using the analytical model (*Δa*=1), using Eq. (S12). **E**: Evolving MGUS progression rates, fitted to data from Rosiñol *et al*. (dots. Non-evolving: 10% at 10 years, 13% at 20 years follow-up. Evolving: 55% at 10 years, 80% at 20 years follow-up) (28), for which we show 95% confidence intervals. Non-evolving MGUS confirms *a* low, constant value of *β* (here 0.007, R^2^=0.996). Evolving MGUS led to a value of *p*=0.071 (R^2^=0.975). **F, G**: Impacts of age at MGUS detection and progression risk reduction *r* on MM-specific mortality as a function of evolving progression rate, calculated using Eq. (S13).

We calculated the cumulative MM-specific mortality following MGUS detection, defined as the probability of an individual to die from MM within a pre-defined number of years after detection of MGUS at a fixed age (27). We distinguished death events from MM and death from other causes. In **Fig. 3B** we display the MM-specific mortality as well as competing risk for MGUS detection ages of 50, 60 and 70 years. In younger groups, the chance to die from MM was comparable to the chance to die due to other causes, the chance to die from other causes increased with age. MM-specific mortality varied strongly with the risk reduction factor *r* (**Fig. 3C**). Equation (S12) proves that MM-specific mortality should not be affected by the screening parameters *a*_0_ and Δ*a*, which only determine the age-specific prevalence of MGUS cases.

### MGUS to MM progression variability and evolving MGUS

Our framework allows to assess the impact of variation in MGUS progression rates (22), as well as the impact of evolving MGUS (28), in which the progression rate changes over time. Variability in MGUS progression rate *p* (per individual per year) can lead to large variability in mortality 10 years post MGUS detection if screening has no effect (*r*=1.0), but this effect is reduced as risk reduction takes effect (*r*<1, **Fig. 3C**).

MGUS patients either belong to a large group of individuals who progress at a constant rate, or to a small group who progress at an accelerating rate (28). Out of 359 MGUS patients reported in (28), 330 (92%) were reported non-evolving and 29 (8%) were evolving (**Fig. 3 E**). We approached these reported progression rates using a discrete-time rate increase, whereby for each individual the rate to progress after exactly *n* years is given by the *β*\*(1−*β*)^n^ (see **Fig. 3**, Supplement). We inferred that nonevolving individuals progress at *β*=0.007, which approximates our constant progression rate well for the majority of individuals of *p*=0.01. In contrasting, evolving MGUS individuals progress with a 10-fold higher value of *β*=0.07. MM-specific mortality increases considerably with evolving MGUS rate (**Fig. 3F**), and decreases with the progression risk reduction (**Fig. 3G**). In addition to population-based variation on progression rates, global migratory effects could make a difference for the impact of MGUS screening. In the Supplement we discuss whether global human migration patterns can have effects on the distribution of genetic variants and the distribution of disease (29). We used data from Ghana (15) as a representative example. Current and realistic levels of immigration of high-risk individuals are very unlikely to impact US MGUS and MM statistics (Fig. S3).

### Equal reduction of MM prevalence can serve as a criterion for optimal screening frequency among high- and low-risk populations

We sought to identify best screening distributions among different risk groups with the goal of minimizing MM prevalence, using Eqs. (S3)-(S8). To illustrate out approach, consider the case of two populations with different lifetime risks of MGUS; in this situation, a fraction *y* of available screenings is applied to the high-risk population. The remainder, 1-*y* of available tests is applied the low-risk population. There exists a value of *y* for which we observe equality of fraction of MM cases in these groups. If *y*=1, all screening effort should be initially directed toward the high-risk individuals. For high values of *r* no value *y* between 0 to 1 can be found (**Fig. 4A**). For the point estimate *r*=0.61 (7), we also found *y*=1 (all screenings should be applied to high-risk). Lower values of *r* could permit values *y*<1 (**Fig. 4B**), ranging from *y*=71% (*r*=0.0) to *y*=96% (*r*=0.3), given *a*_0_=50y and being less sensitive to Δ*a* (**Fig. 4C**, **and Table S5**; *y* was between 81-93% for Δ*a*=1 and between 79-95% for Δ*a*=4 (fixed *r*=0.1, **Fig. 4D**, Table S6).

**Figure 4:**
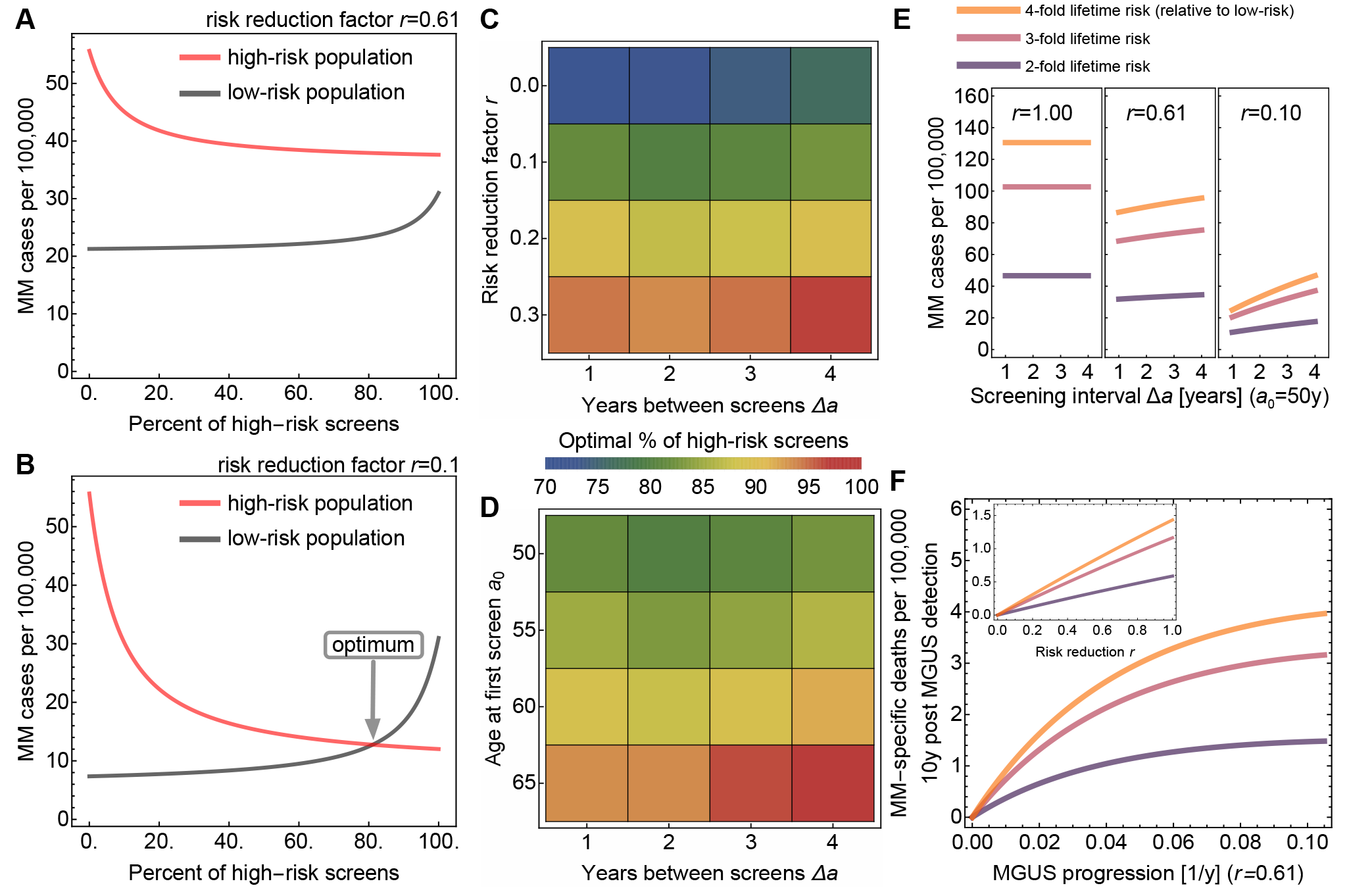
Equal disease fractions as a criterion for optimal screening distribution. **A, B**: Comparing MM fractions in the high-risk and the low-risk populations (males and females, respectively), with *a*_0_=50, *Δa*=1y, for different *r*. For *r*=0.61, equality could not be observed for any percentage of high-risk screens (**A**). For *r*=0.1, equality was observed at about 81% high-risk screens (**B**). Thus, an optimal fraction of screens was defined as the point where the fractions of MM cases in both sub-populations were the same. **C**: Location of the optimal fraction (see scale) under variation of *r* and Δ*a* (see Table S5), *a*_0_=50y. Changing *r* from 0 to 0.3 would lead to up to 20% change in the optimal high-risk fraction of screens. Changing Δ*a* from 1 to 4 would lead to 1-3% change in the optimal high-risk fraction of screens. **D**: For fixed *r*=0.1, changes in *a*_0_ had more drastic effects than changes in Δ*a*, (also Table S6). **E**: For risk groups with a lifetime risk is higher that 2-fold, we examined the effect of risk reduction and screening interval (*a*_0_=50) on the number of MM cases (Eq. S8). **F**: MM-specific deaths per 100,00 were calculated as the product of screened MGUS individuals at age 60 and the 10-year follow-up MM-specific mortality (*a*_0_=50, Δ*a*=1, age at MGUS detection 60y). Both risk reduction and spacing of screens have more pronounced effects in higher risk groups.

Lifetime risk of MGUS affected the optimal Δ*a* for a high-risk population, when Δ*a* was fixed for the low-risk population (**Fig. S3B**). Low-risk screening every 10 years would lead to equal prevalence reduction if high-risk screening was implemented every 3 years provided high-risk individuals experience a 2-fold lifetime risk. If this risk were 4-fold, the ratio would be 10:1 (**Fig. S3C**).

### Higher than 2-fold lifetime risk-groups could benefit strongly from regular screening

Multiple factors that determine increased lifetime risk of MGUS, notably family history of MM (30). We analyzed the sensitivity of MM prevalence and MM-specific mortality to screening frequency and risk reduction. Both risk reduction and spacing of screens have more pronounced effects in higher risk groups, but in those groups steeper increase in mortality was observed with decreasing screening frequency (**Fig 4E**). Importantly, the increase in MM-specific deaths saturated with increasing progression rate, indicating that in high-risk groups, mortality reduction could be achieved in subgroups of intermediate progression rates (**Fig 4F**).

## SUMMARY AND CONCLUSIONS

Multiple myeloma (MM) remains incurable for the majority of patients, and decreasing mortality is of as much interest as decreasing its prevalence (10). All patients appear to progress to symptomatic MM from a pre-malignant, asymptomatic stage called MGUS (31). There are outstanding diagnostic tests for MGUS, which implies the possibility of delaying progression of MGUS to MM by screening and early identification (32). We examined the stochastic onset and progression of MGUS in a population model of high-and low-risk individuals. We evaluated a range of possible screening strategies based on the consideration that diagnosis of MGUS permits progression reduction due to several possible interventions or modifiable risk factors such as aspirin, metformin, or mediation such as exercise or diet alterations in obese MGUS patients (7, 8, 10, 11, 32).

Our approach allowed us to quantify the amount of risk reduction needed to result in certain reductions in MM-specific mortality and MM prevalence (measured as MM fraction). To avoid lead-time bias, we evaluated screening scenarios in terms of mortality and MM prevalence. Length-time bias, on the other hand, is a form of selection bias that occurs due to heterogeneity in progression speed of a malignancy. This bias was absent in our study as we modeled uniform progression of the disease, i.e. a high-risk person with early incidence of MGUS progressed equally fast to MM as a low-risk person with late MGUS incidence; the time spent in the MGUS state in the no-screening scenario was independent of age (13). Therefore, these common sources of bias in epidemiologic prevention studies did not confound our results.

Using a stochastic simulation framework and an analytical model we measured MGUS and MM prevalence and MM-specific mortality in different risk-groups, for different screening strategies and varying progression risk reduction after MGUS detection. For effective MM prevalence reduction, better screening results are expected for early as possible screening and frequent follow-up. Improved chemoprevention, effectively reducing progression risk, may also reduce MM-specific mortality. We found that this effect is more pronounced in individuals with evolving MGUS, and especially in individuals with higher than 2-fold lifetime MGUS risk.

A range of screening scenarios can be studied with our theoretical framework; this approach allows us to evaluate how screening parameters influence MM prevalence and MM-specific mortality. We did not explicity address screeing toxicity here, nor did we model smoldering multiple myeloma (sMM)—an intermediate stage between MGUS and MM with a much higher rate of progression to full MM of about 30% per year— in part because it remains unclear whether sMM is a requisite intermediate between MGUS and MM. Our framework can be adjusted and expanded. It might become useful to include screening strategies that can identify sMM early. This expansion could reveal that treating only high-progression-risk sMM cases might be most effective.

Assessments of screening and prevention in solid tumors, e.g. prostate cancer, have been controversial, and lacking of evidence for screening in large prospective trials (33). We share skepticism of potential "medicalization" of asymptomatic conditions. However, the biology of MGUS and the robust laboratory tests demand careful evaluation of the role of screening and prevention. With notable similarities in the epidemiology of prostate cancer and MGUS—most low-grade lesions will not proceed to lethal disease—major differences in technology of screening tests for these diseases are critical. PSA tests for prostate cancer are burdened by substantial false positive (21 - 32% sensitivity) and false negative rates (85 - 91% specificity) (34). In contrast, serum testing for MGUS is straightforward. The sensitivity of SPEP and FLC testing for MGUS is close to 100% and specificity is 99% (35). These differences underline the evaluation of a role of screening and prevention in MGUS/MM. We have shown that the reduction of MM cases and MM specific mortality in high-and low-risk sub-populations could be achieved, but only for drastic reduction in progression risk. Until highly effective agents are developed, identification and follow-up of high-risk individuals are important. Screening for MGUS may have significant population benefits by lowering the incidence of MM, provided effective and non-toxic interventions can be identified. Without further study of chemoprevention strategies, regular screening of MGUS candidates should start as early as possible, with bi-annual follow-up, and focus on high-risk individuals especially with a family history of MM or on groups with strong indication for evolving MGUS progression.

## ACKNOWLEDGEMENTS

The authors gratefully acknowledge feedback by Graham Colditz (St Louis) and Nicola Camp (Salt Lake City), as well as by members of the Michor lab at the Dana-Farber Cancer Institute.

## AUTHORSHIP CONTRIBUTIONS

P.M.A., T.G., M.H.T., and F.M. conceived the study. P.M.A. and J.F. performed computational experiments. PMA performed mathematical modeling and statistical analyses. P.M.A., J.F., and F.M. analyzed the data. P.M.A., T.G., M.H.T., and F.M. wrote the manuscript. P.M.A., T.G., M.H.T., and F.M. supervised the project.

